# Agent-driven Model Development for RNA 3D Structure Prediction

**DOI:** 10.64898/2026.09.08.749228

**Authors:** Maciej Majewski, Laura Malo, Ariadna Montero-Blay, Matteo Marengo, Ricardo Nascimento Dos Santos, Paraskevi Gkeka, Hervé Minoux

## Abstract

Large language model (LLM) agents have shown promise in driving scientific discovery, but their effectiveness in complex, real-world biological problems remains underexplored. We ask whether a general-purpose LLM agent can drive semi-autonomous development of a model for a genuinely hard biological problem, RNA 3D structure prediction. We designed a development loop where, under a fixed budget and with human supervision, the agent iteratively proposed, implemented, trained, and evaluated model changes. Over 297 iterations, the model evolved from a randomly-initialised baseline to QuickFold, an 8.9 M-parameter folding trunk that matches the strongest open-source baselines (RhoFold+, NuFold) on lDDT and TM-score within noise on a held-out test set at a fraction of their inference cost. We frame this less as a new predictor than as a case study in feedback-driven, agent-led model development.

## 1 Introduction

Predicting RNA three-dimensional structure from sequence is among the most challenging open problems in structural biology. RNA folding is governed by conformational flexibility, non-canonical base pairing, and pseudoknots Schneider et al. (2023), and the field is constrained by a severe data imbalance: only ~10,000 RNA structures have been deposited in the Protein Data Bank (PDB) against millions of known sequences Bernard et al. (2024); rna (2026). The emergence of deep learning, accelerated by AlphaFold2’s success on proteins Jumper et al. (2021), has driven development of RNA-specific methods such as trRosettaRNA Wang et al. (2023), RhoFold+ Shen et al. (2024), and NuFold Kagaya et al. (2025), substantially improving accuracy on benchmark families. Yet performance degrades on non-canonical, non-ribosomal, and novel RNAs, demanding continued, iterative model development across architectures and featurizations Schneider et al. (2023); Das et al. (2023).

Large language models (LLMs) have recently emerged as drivers of such iterative development. Early systems, the AI Scientist Lu et al. (2024), Google DeepMind’s AI Co-Scientist Gottweis et al. (2026), and Kosmos Mitchener et al. (2025), demonstrate end-to-end automation of hypothesis generation, experimentation, and synthesis across scientific domains. A more constrained but fully closed-loop instantiation is autoresearch Karpathy (2026), in which an agent repeatedly edits a training script, runs a time-boxed experiment, and keeps or discards changes by measured performance. Prior closed-loop systems, however, were mostly shown on toy ML benchmarks or on tasks with a small search space and a cheaply computed, fixed metric. RNA 3D structure prediction is harder on both fronts: each iteration is costly to evaluate, and the optimisation target itself shifted during the project, as structural-validity terms had to be added mid-run. This paradigm has been extended to agentic neural architecture search Jeong et al. (2026) and to drug discovery, where DrugSAGE Zhang et al. (2026) and closed-loop molecular property prediction Ning et al. (2026) show that agentic loops can discover generalizable modeling improvements.

Motivated by these developments, we ask whether an LLM agent, operating under human supervision, can drive model development for a genuinely complex biological task such as RNA 3D structure prediction. We let the agent drive the development of a new model while we steer it through fixed objectives, requirements, and feedback, rather than by designing the architecture ourselves. The resulting model, QuickFold, was evaluated and compared against similar published tools. Our contributions are threefold: (i) a semi-autonomous, feedback-driven workflow adapting the autoresearch loop to a hard biological problem; (ii) QuickFold, a compact model matching open-source RNA predictors at a fraction of their inference cost; and (iii) an analysis of the human–agent division of labour that made this possible.

## 2 Agentic workflow

To test whether an agent can successfully drive model development for a biologically relevant case, we picked a specific problem: build a model that predicts RNA 3D structure based on its sequence. The designed workflow draws inspiration from the autoresearch repository (Karpathy, 2026) and automates the actions a machine-learning (ML) scientist routinely takes during a model development. It is realised entirely inside the Cursor harness as a set of rule files, prompts, and subagents driving a single LLM (Claude Opus 4.8). The workflow ran under a deliberately tight compute budget on a single machine with one GPU (L40S, 46 GB). The workflow has two phases (Fig. 1): a human-supervised *setup* that produces a working baseline, and an *autonomous experimentation loop* that improves it.

**Figure 1.**
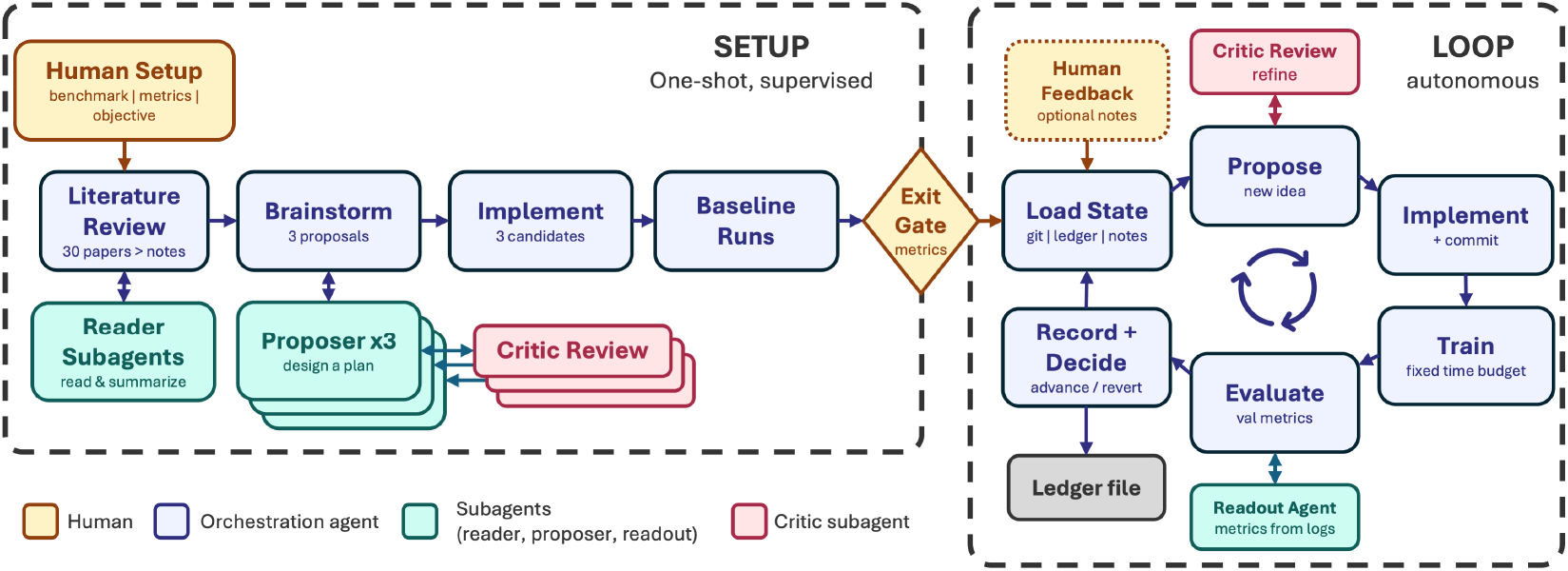
Agentic workflow. A one-shot, human-supervised setup phase (literature review, brain-storming with proposer and critic subagents, implementation of three candidates, baseline selection) passes through an exit gate into an autonomous experimentation loop (load state, propose one change with critic, implement and commit, train, evaluate, record, and decide). Colour distinguishes human, agent, and subagent actions.

### 2.1 Setup

The setup stage is supervised by the user and runs once. The human prepares the materials the agent is not trusted to gather and fixes the ground rules. Specifically, the human: 1) collects the relevant literature and tools (≈30 papers); we deliberately provide this corpus to prevent abstract-only hallucination on paywalled sources and to hold the knowledge base constant across runs; 2) curates and freezes the benchmark (a set of single-chain RNAs, 16–500 nucleotides: train 18,675 / val 68 / test 80, split temporally by deposition date so the test set (deposited from 01/01/2024 onward) post-dates the training cutoffs of the reference tools; SI.A.1); 3) defines the metrics (lDDT, TM-score, eRMSD, and clash rate; SI.A.2) and the optimization criterion (primary: validation lDDT); and 4) sets the run constraints: training time budget, inputs mandatory for this run, and all models trained from scratch with no fine-tuning. After that, the user kicks off the agent part of the setup.

Given these materials, the agent reads and summarizes every paper through reader subagents, making a summary note for each article that will be accessible to other agents. The agent then brainstorms at least three distinct candidate architectures via proposer subagents refined by a critic subagent (the roles of the main agent and each subagent are described in SI.A.3), then implements, trains, and evaluates each design and selects a baseline by the set objective. The phase ends at an exit gate where the human reviews and accepts the pipeline, outputs, and baseline before the loop may begin.

### 2.2 Experimentation loop

The accepted baseline seeds the loop, which is designed to run with minimal supervision. Each iteration is a fixed cycle: load state (git status, ledger, current best) → propose one change, vetted by the `critic` subagent → implement and commit → train within a fixed time budget → evaluate on the validation set → record to an append-only ledger and decide. A change is accepted only if validation lDDT improves over the current best by more than 0.001; otherwise the commit is reverted and the agent tries something else.

The training budget was chosen to maximize the number of iterations rather than the quality of any single one. It began at 10 min and was progressively raised to 8 h as returns from cheap runs were exhausted; lifting it proved one of the most effective performance levers. Early on, agent reasoning and GPU training ran sequentially, idling the GPU; from iteration ~200 the agent instead planned several iterations at once and submitted batched overnight trainings (20–24 h budget), with the human triggering analysis about once a day.

Two rules kept the agent running. First, a “never stop” rule forbids the agent from pausing to ask whether it should continue. Second, an “escape-local-minima” rule: after three consecutive non-improving experiments, abandon hyper-parameter tweaks in favor of radical architectural changes. This rule the agent cites as the trigger for its two largest architectural moves.

Human feedback is optional and enters the loop through an append-only `Notes.md` that the agent reads at the beginning of each iteration. This feedback derives from the human inspecting the outputs (e.g., visual inspection in a molecular viewer), signaling the issues that no scalar metric captured. When the agent stalls it is prompted to brainstorm and log ideas to a `Brainstorm.md` file for use in the current or a later iteration.

The resulting division of labour is a central observation: the *agent* owned the metrics, mechanisms, and implementation, while the *human* owned visual structural judgement, and the scope of the work, steering the trajectory without explicitly designing the architecture.

## 3 Results

### 3.1 Model evolution

We attempted many optimization runs (SI.A.4). In this specific run, the setup was: Claude Opus 4.8 as the agent LLM, validation set lDDT as the optimization objective, and the sequence and precomputed shallow multiple-sequence alignments (MSAs; generation and depth in SI.A.1) as the inputs. The setup phase returned three candidate baselines that differed in their input features (language-model, MSA, or a hybrid). The hybrid approach (`lm_msa`) was selected at validation lDDT 0.393 and seeded the loop.

Over 297 iterations spanning roughly one month, the agent tried many changes rapidly. Most were dead ends that were systematically logged and reverted, while a minority advanced the model, which grew in architectural complexity, parameter count, and training time. The effect of the model changing on a single target is visible in the predictions of `8a3d_B` (Fig. 2a). Progress was *discontinuous*: a handful of accepted iterations separated by long flat stretches of failed trials (green vs. gray points, Fig. 2b). The largest gains came from: a ViennaRNA Lorenz et al. (2011) secondary-structure pair bias (event 2, Fig. 2), an exponential moving average (EMA) of the weights (event 3, Fig. 2), a corrected nucleotide template (a bug fix; event 5, Fig. 2), an AlphaFold2-style triangle-multiplicative pair update (event 6, Fig. 2), and raising the training budget from 20 → 480 min together with a crop curriculum.

**Figure 2.**
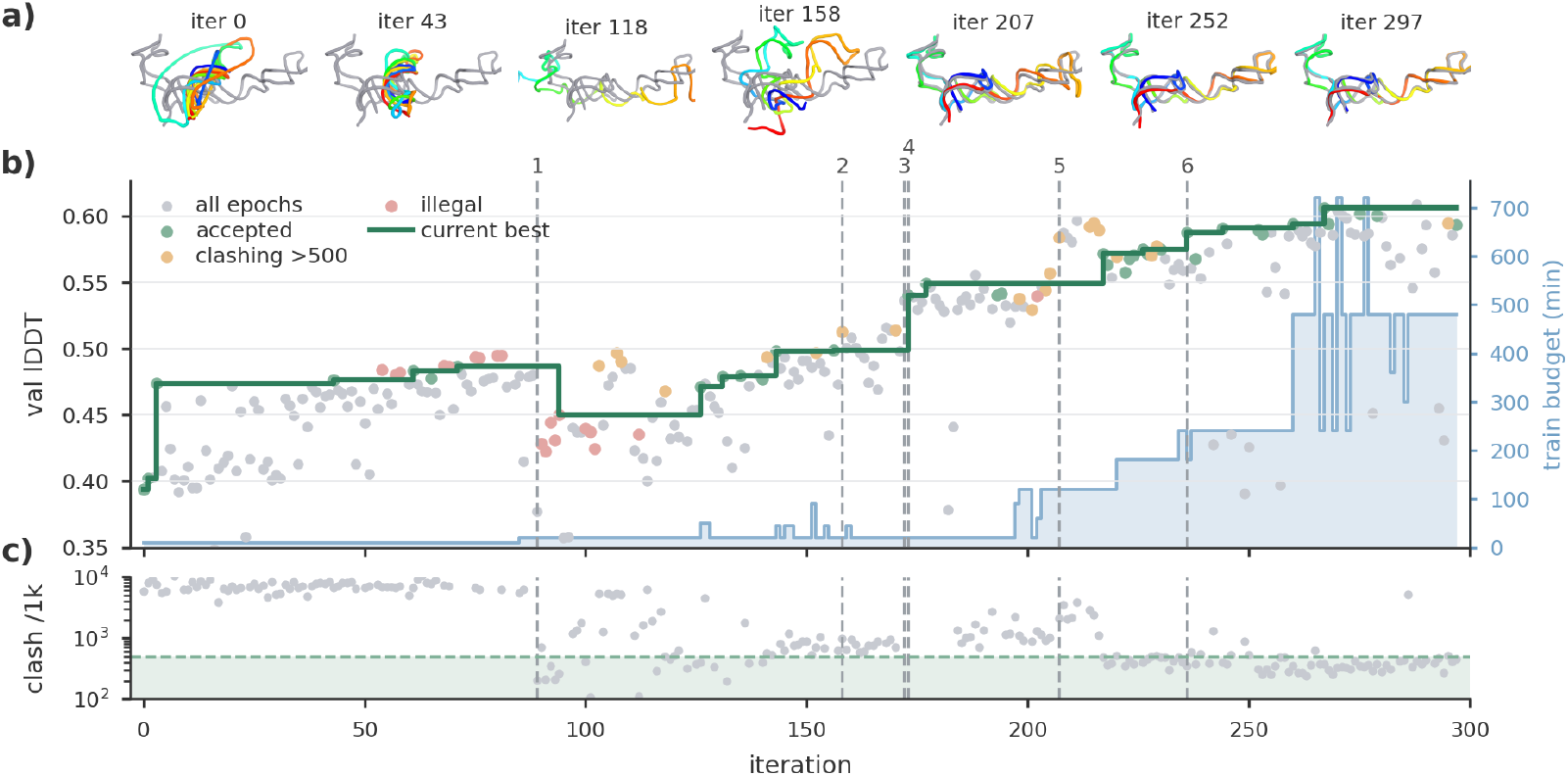
Model training trajectory over 297 iterations. (a) Cartoon renders of the predicted structure (rainbow) overlaid on the native reference (gray) at seven representative iterations (PDBid: `8a3d_B`). (b) Validation lDDT versus iteration. Gray, all epochs; green, accepted models; orange, accepted models whose clash rate exceeds the acceptable threshold (>500 per 1k) once structural correctness was enforced (1); red, illegitimate “advances” excluded from selection (ensemble or fine-tuned models). The dark-green step line is the current best legitimate, non-clashing model; it climbs early on, after non-clashing structures became a hard requirement (1), drops at iteration 94 to the best genuinely non-clashing structure (lDDT ≈ 0.45), and recovers thereafter. The blue shaded area (right axis) is the per-iteration training budget (minutes). (c) Reported clash rate (clashes per 1,000 heavy atoms, log scale, capped at 10^4^); the green band marks the acceptable region (≤500 per 1k). Numbered dashed lines mark methodological milestones: **1** structural correctness required (iter. 89); **2** secondary-structure prediction (158); **3** EMA of weights (172); **4** model selection objective changed to hybrid TM-score / lDDT (173); **5** clash-metric and nucleotide-template bug fix (207); **6** triangle-multiplicative pair update (236).

Two human-imposed changes to what counted as an acceptable model reshaped the trajectory. First, requiring geometrically valid, clash-free predictions (iter. 89; event 1) rejected several lDDT-improved but clashing models (orange points, Fig. 2b) and drove the clash rate down by over an order of magnitude (Fig. 2c). Second, at iteration 173 (event 4) the selection objective became an equally weighted TM-score/lDDT average, so local gains could no longer come at the cost of the global fold. A separate fix corrected a faulty clash metric (event 5, iter. 207) whose blind spot had masked collapsed geometry, prompting the template bug fix above.

The evolving objective grew the loss from an lDDT-centric frame-aligned point error (FAPE) + distogram + confidence set into a ten-term multi-stage loss (SI.A.6.2); some terms were proposed by the agent, others imposed by the human for structural validity. The governing principle was *scheduling*: geometric and global-fold penalties destroy an unformed fold, so they were warmed up late in training.

Human feedback also drove the secondary-structure prior, repeated budget extensions, and finally dropping the MSA; the language model made it unnecessary at inference, and the final MSA-free variant (iter. 297) matches the MSA model within seed noise.

### 3.2 Final model

Over almost 300 iterations the loop produced QuickFold, a ~8.9 M-parameter folding model with a non-trivial architecture. A frozen RNA language model (RNA-FM Shen et al. (2024), as in RhoFold+) supplies sequence embeddings and a ViennaRNA base-pair probability matrix supplies a secondary-structure bias (secondary-structure information is likewise used by NuFold). Together these condition a compact ten-block trunk with an AF2-style triangle-multiplicative pair update, followed by an invariant point attention (IPA)-style structure module that places all heavy atoms by applying predicted rigid frames to idealized per-nucleotide templates. The individual modules are thus known and explored in other tools; the novelty is, however, their arrangement, and the assembly is deliberately efficient: the 8.9 M-parameter trained trunk compares against 27.2 M for RhoFold+ and 94.8 M for NuFold (SI, Table 4). A detailed description and an architecture diagram are given in SI.A.6.1.

On the held-out test set (*n* = 80), QuickFold matches RhoFold+ and NuFold on lDDT and TM-score within seed noise while being the fastest by a wide margin (Table 1): its smaller trunk and MSA-free design fold a target in ~3.5 s, against ~20 s for NuFold and ~275 s for RhoFold+, with inference timed excluding MSA search. Seed noise is small: across three seeds QuickFold reproduces at test lDDT 0.600 ± 0.016, which sets the resolution below which single-run differences are not meaningful.

**Table 1.** Held-out test benchmark against open-source methods (*n* = 80). *Mean of 3 predictions* averages over three random seeds for QuickFold and the three ranked predictions for NuFold (standard deviation in parentheses); RhoFold+ is deterministic, so its single prediction is repeated. *Most confident* is the prediction chosen by predicted confidence (QuickFold: pLDDT; NuFold: top-ranked; RhoFold+: the single prediction). Inference time is average per target and excludes MSA search. Bold marks the best value per accuracy column.

| Method | Mean of 3 predictions |  |  |  | Most confident |  |  |  | time (s)↓ |
| --- | --- | --- | --- | --- | --- | --- | --- | --- | --- |
|  | IDDT↑ | TM↑ | eRMSD↓ | clash/1k | IDDT↑ | TM↑ | eRMSD↓ | clash/1k |  |
| **QuickFold** | **0.600** (0.016) | **0.306** (0.030) | 1.638 (0.032) | 160.4 (12.1) | **0.613** | **0.324** | 1.600 | 140.9 | **3.5** |
| NuFold | 0.591 (0.002) | 0.283 (0.001) | **1.396** (0.011) | 93.6 (8.0) | 0.593 | 0.284 | **1.386** | 87.1 | 20.0 |
| RhoFold+ | 0.531 | 0.272 | 1.653 | 1424.8 | 0.531 | 0.272 | 1.653 | 1424.8 | 274.9 |

## 4 Discussion

The workflow enables rapid prototyping rather than fully autonomous research, yet still compresses model development from months to weeks. It introduces a feedback-driven mode of work: the human neither pre-plans nor enforces an architecture; the agent writes the code, brainstorms alternatives, and keeps whatever improves the metric, while discarded ideas survive only as a ledger entry. Human input is most useful as feedback on concrete failures or as a change of requirements (SI.A.8, Table 8).

The timing of that input matters. Early feedback can over-constrain the evolving model, whereas feedback given once predictions already look reasonable is far more productive. The loop was intended to be autonomous but in practice halted occasionally when the agent judged itself stuck.

When the agent stalls, even a generic “be bolder” encouragement often restarts progress. Of the 98 human turns, roughly 60 were variants of this “be bolder” template; the remainder carried concrete input: scientific suggestions from visual inspection, bug reports, resource authorisation, and scope decisions (SI.A.8). The compute budget is itself a design choice: fixing a per-iteration training time shifts the objective from “the best model” to “the best model trainable under that budget,” which biases the search toward a smaller, faster-inference architecture.

Two practical caveats temper the results. First, small validation sets make many single-run improvements indistinguishable from seed noise (measured at *<*0.01 lDDT); several mid-campaign “wins” were later reinterpreted as noise, so later and more modest changes should be judged on repeated runs and averages rather than single deltas. Second, an autonomous agent generates an unmanageable volume of code, and the final model must be cleaned and carefully inspected before any release.

Finally, the honest limitations. Global fold remains the weak axis (test TM ≈0.31): although QuickFold leads the reference methods on this metric, its absolute quality is still modest. The ≈0.60 lDDT plateau appears to be a data ceiling rather than an architectural one. This style of development thus makes it comparatively easy to reach the level of existing methods but hard to push meaningfully beyond them. Its value is twofold: it lets ML scientists explore more options than they otherwise could, and it lets domain experts without deep architectural knowledge steer model building through feedback and prior knowledge.

In sum, a general-purpose LLM agent, held to a fixed budget and steered by human feedback rather than human design, drove the development of QuickFold from a random initializer to a compact model competitive with established RNA structure predictors. The contribution is less the artefact than the demonstration that feedback-driven, agent-led development is a practical mode of building models for hard scientific problems; one that widens who can build them and how many ideas can be tried, even if pushing past the current state of the art remains the harder, still-human problem.

## A. Supporting information

### A.1 Benchmark curation

The benchmark is a curated set of single-chain RNAs derived from the Protein Data Bank (PDB): train 18,675 / val 68 / test 80 sequences, with lengths of 16–500 nucleotides (median ~84 nt).

#### Data split

Chains were split by PDB deposition date so that later structures are never seen in training: *train* deposited on or before 2021-12-31, *val* between 2022-01-01 and 2023-12-31, and *test* on or after 2024-01-01. The test cutoff (2024-01-01) is deliberately more recent than the training-data cutoffs of both reference tools (RhoFold+ and NuFold), so the held-out test set post-dates the data those baselines were trained on and is unseen by QuickFold and the baselines alike. The test-set PDB identifiers are listed in Table 2.

**Table 2.**
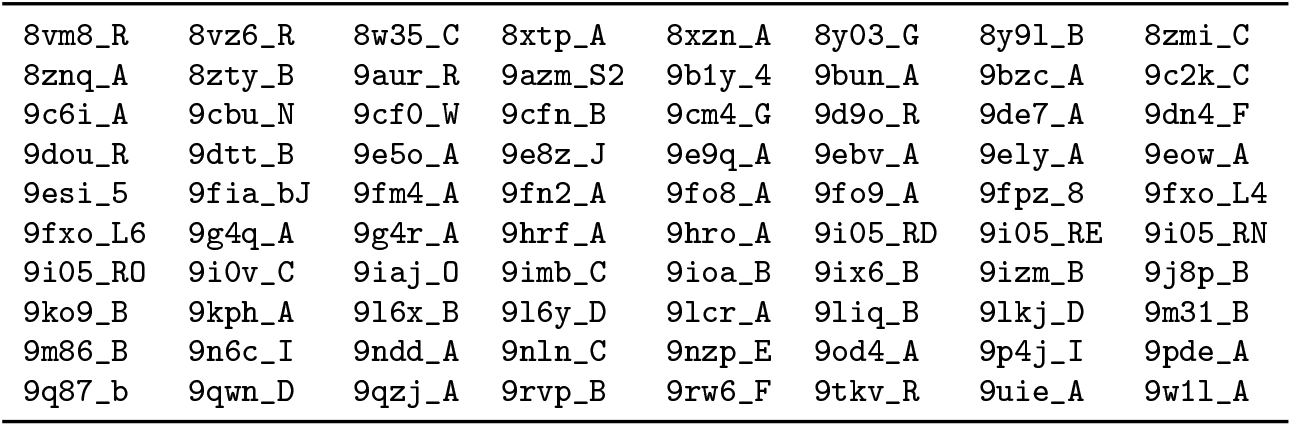
Held-out test set (*n* = 80): PDB identifier and chain for each target (format pdbid_chain). All entries were deposited on or after 2024-01-01.

#### Filtering and redundancy reduction

Starting from RNA-containing PDB entries (X-ray, cryo-EM, and NMR; resolution 5 ≤ Å), we retained single chains of 16–500 nucleotides with limited missing density and few non-standard residues; modified nucleotides were mutated to their standard parents where the assignment was unambiguous. Sequences were made non-redundant with a graduated mmseqs2 clustering cascade (sequence-identity levels 0.5–0.95 at 0.8 coverage), keeping one high-quality representative per cluster. A conformer-aware step additionally preserved sequence-identical but structurally divergent copies (pairwise USalign TM-score *<*0.45) as separate representatives, so that distinct folds of the same sequence are not collapsed.

#### Chain fragmentation

To enlarge the training pool, training chains were fragmented at secondary-structure junctions. Using an automatic secondary-structure annotation — the `annotator` CLI of the RNApolis package Zok (2024), run in extended mode (`-e`), which derives Leontis–Westhof-classified base pairs (canonical, non-canonical, and pseudoknotted) directly from the deposited 3D coordinates rather than from a thermodynamic prediction — candidate cuts were placed only in single-stranded regions between stems (never inside a base-paired stem), and contiguous unions of the resulting structural blocks, together with individual stem-loops, were emitted as sub-fragments of at least 16 nt. Validation and test sets were *not* fragmented: they contain only intact full single-chains.

#### Multiple-sequence alignments

Both QuickFold’s original `lm_msa` inputs and the alignments supplied to the RhoFold+ and NuFold baselines were precomputed once and frozen. Alignments were built with RhoFold+’s single-pass `blastn` search (NCBI BLAST+, `-task blastn`, `-evalue 0.001`) against a full local copy of RNAcentral rna (2026), converted to A3M through the bundled rMSA-derived scripts; no iterative profile expansion or covariance-model search was run. The resulting alignments are shallow for most targets: 29/80 test targets are query-only (depth 1), the median depth is 3, and only 15/80 reach RhoFold+’s 128-row cap. This depth is a genuine property of RNA homology search under our compute budget rather than a truncation artefact, and the MSA-conditioned baselines are expected to underperform their typical accuracy on this shallow-alignment regime (SI.A.7, Table 7). QuickFold’s deployed variant is MSA-free and is unaffected.

### A.2 Metrics definition

All metrics compare a predicted structure to its experimental reference over all heavy atoms (hydrogens ignored), matching atoms by residue index and atom name. The optimization criterion throughout was the primary validation set lDDT.

#### lDDT

The local Distance Difference Test scores Mariani et al. (2013), without superposition, the fraction of reference interatomic distances that are preserved in the prediction. For every pair of reference atoms closer than a 15 Å inclusion radius, the distance is counted as preserved if it is reproduced within one of four tolerances (0.5, 1, 2, 4 Å); the per-atom score averages over the four tolerances and the global lDDT averages over all atoms with at least one neighbour. Higher is better.

#### TM-score

The “TM” column of Table 1 is the TM-score computed by USalign Zhang et al. (2022) in RNA mode with an optimal residue alignment, anchored on the C3^*′*^ atom and normalised by the reference length. Values lie in [0, 1]; higher is better. It captures global fold quality, complementing the local lDDT.

#### eRMSD

The eRMSD is an alignment-free, base-orientation-aware measure of nucleobase arrangement (dimensionless, lower is better), computed with `barnaba` at a 2.4 cutoff Bottaro et al. (2019).

#### Clash rate

We report a MolProbity-style clashscore: two heavy atoms clash when their van der Waals radii overlap by more than a 0.4 Å tolerance, after excluding true covalent (1–2 and 1–3) neighbours and the phosphodiester linker atoms; the score is the number of clashing pairs per 1,000 heavy atoms. Native structures, measured with the same clashscore over the 80 native chains of our test set, score ≈18 per 1k.

#### Extended evaluation

For the detailed results (Table 5) we additionally report a global-alignment TM-score (TM_glob_) and a root-mean-square deviation (RMSD), both computed with `barnaba` on C1^*′*^ atoms after a single global least-squares (Kabsch) superposition of the whole structure. RMSD is the root-mean-square distance (in Å, lower is better) between corresponding C1^*′*^ atoms after that superposition. These global measures complement the local ones: lDDT and the main-text TM-score (TM_loc_) use USalign’s optimal residue alignment, which superposes the best-matching residue core and so rewards well-modelled local regions, whereas the barnaba global alignment fits all residues at once and is correspondingly more sensitive to errors in the overall fold.

### A.3 Agent roles and subagents

The workflow is driven by a single LLM (Claude Opus 4.8) that adopts several roles, realised as prompt-scoped subagents within the Cursor harness (Fig. 1). Colour in that figure distinguishes the human, the main agent, and the subagents.

- **Main agent** — the orchestrator that executes the iteration cycle: it loads state, edits and commits code, launches training and evaluation, and writes the ledger. It owns the control flow and delegates focused tasks to the subagents below.
- **Reader subagents** — during setup, each reads one paper or tool from the fixed knowledge base and writes a structured summary note the other agents can consult, so the literature is digested once and shared.
- **Proposer subagents** — brainstorm candidate architectures or changes, each contributing a distinct design so the search does not collapse onto a single idea.
- **Critic subagent** — reviews and vets each proposal before implementation, checking it against the objective, the run constraints, and past ledger entries, and flags weak or redundant ideas.
- **Readout subagent** — parses the raw training and evaluation outputs (metrics and logs) into the structured ledger entry on which the accept/reject decision is made.

### A.4 Other optimization runs

Beyond the run reported above, we carried out additional optimization runs that varied the initial conditions along four axes: the primary objective (lDDT or TM-score; the reported run used lDDT), the input (sequence and MSA-based or single-sequence), the agent LLM (Claude Opus 4.7, Claude Opus 4.8, or GPT 5.5), and whether structural correctness (bond and angle validity) was enforced from the start or only introduced later in the run. Three of these runs reached the performance level of the reference models.

Several observations recurred across runs. First, over-constraining the search early was harmful: runs that enforced geometric validity from the beginning tended to stagnate, producing structurally valid but trivial folds that subsequent iterations could not escape. Introducing the validity requirement only after a rough fold had formed avoided this trap, consistent with the late-warmup scheduling of SI.A.6.2. Second, successful runs tended to converge on a shared set of essential components; in particular, secondary-structure prediction emerged as an intermediate in *all* successful runs.

These runs also underscored the importance of specifying the task precisely. In one run the objective was stated only as sequence in, 3D structure out; the model satisfied this by predicting a single atom per nucleotide and omitting the remaining atoms. The requirement was too loose - it was corrected mid-run for that campaign and tightened (an explicit all-atom output) for subsequent runs.

### A.5 Detailed model evolution

#### Phases and groups of solutions

The 297 iterations fall into a sequence of exploration phases rather than a smooth climb. Early iterations (0–90) established deep-supervision frame-aligned point error (FAPE) and annealed the initial-frame noise; this phase was throughput-rather than capacity-limited, so the agent favoured cheaper crops and short budgets. A middle phase (90–200) added the ingredients that made structures usable: a connected-chain decoder, a differentiable soft-lDDT loss, the ViennaRNA Lorenz et al. (2011) base-pair-probability bias (iter. 158, the largest raw-lDDT jump of this era), an exponential moving average (EMA) of the weights (iter. 172), and a crop curriculum with a longer budget. A late phase (200–279) raised capacity once training was no longer the bottleneck: the corrected nucleotide template (iter. 207, the single largest gain), a glycosidic-*χ* head, late-warmup validity terms, and the AlphaFold2 (AF2) triangle-multiplicative pair update (iter. 236), the first change to beat the standing best on every axis. A final phase (280–297) confirmed that RNA-FM was the right frozen encoder (a sweep of seven RNA language models found no better) and that the multiple-sequence alignment (MSA) could be removed at inference essentially for free (iter. 297).

#### Loss-function evolution

The training objective grew with the model, from an lDDT-centric set (FAPE + distogram + predicted lDDT [pLDDT]/predicted aligned error [PAE]) to a ten-term objective as requirements expanded: soft-lDDT (metric-aligned) → Kabsch soft-TM (once TM became a 50/50 objective) → late-warmup validity terms (van der Waals clash, backbone continuity, bond angles) → a glycosidic-*χ* term. Some terms were proposed by the agent, others imposed by the human to enforce structural validity. The governing principle was *scheduling*: geometric and global-fold penalties destroy an unformed fold and help only once one exists, so they were warmed up late in training (past roughly 60% of the wall-clock budget).

### A.6 Final model

#### A.6.1 Model architecture

QuickFold (iter. 297 variant) is an MSA-free model with an 8.9 M-parameter trained trunk (single-representation width *d*_*s*_ = 256, pair width *d*_*z*_ = 128). The forward pass runs sequence → featurizer → ten trunk blocks → a four-iteration structure module → heads and atom placement, with one recycle (two full passes at inference); Fig. 3 summarises the flow.

**Figure 3.**
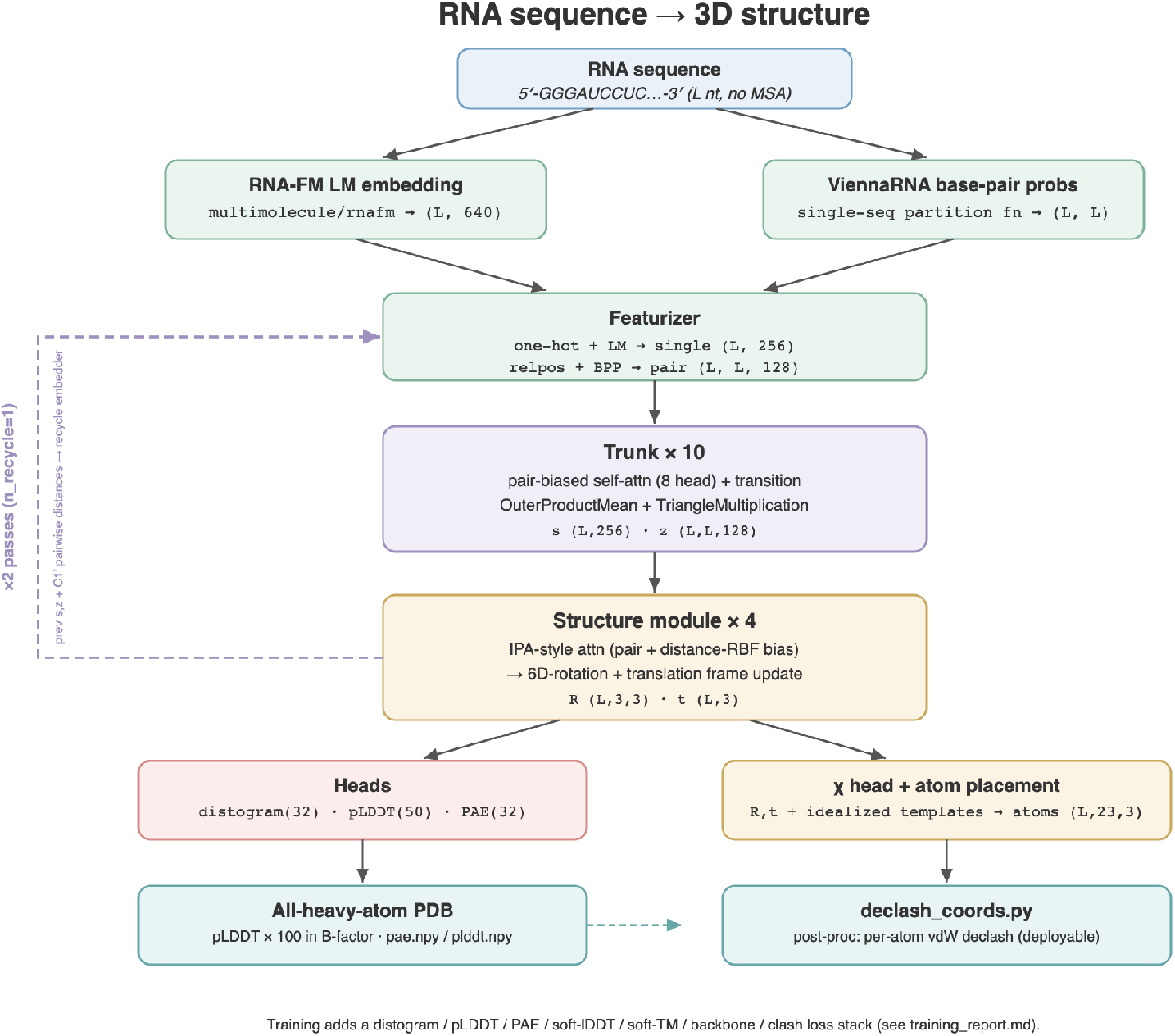
Simplified QuickFold architecture (deployed MSA-free iter. 297). Frozen RNA-FM embeddings and ViennaRNA base-pair probabilities feed a ten-block trunk with triangle-multiplicative pair updates and an invariant point attention (IPA)-style structure module; heavy atoms are placed from predicted frames and idealised per-nucleotide templates, with a post-hoc declash on the deployable path.

##### Inputs

Two frozen, single-sequence signals condition the network. A frozen RNA-FM language model Shen et al. (2024) provides per-nucleotide embeddings ((*L*, 640)) projected into the single representation — the same class of feature used by RhoFold+. ViennaRNA base-pair probabilities provide a secondary-structure signal ((*L, L*)), injected as a pair feature through a zero-initialised, bias-free linear map, so that the ~92% of entries with negligible pairing probability contribute exactly zero and the bias fires only on confident base pairs.

##### Trunk and structure module

Each trunk block applies pair-biased self-attention on the single representation with a transition, an OuterProductMean, and a gated (outgoing and incoming) triangle-multiplicative pair update imported from AlphaFold2. The structure module is an IPA-style stack that maintains a per-residue rigid frame, updated through a rotation-plus-translation parameterisation.

##### Atom placement and heads

Heavy atoms are never regressed directly. The predicted rigid frame is applied to an idealised per-nucleotide template, and a small zero-initialised *χ* head rotates only the base-ring atoms about the glycosidic bond; intra-residue geometry is thus chemically valid by construction, and all learned error lives in the inter-residue frame and the *χ* rotation. The idealised templates are not synthetic geometries: for each of the four canonical nucleotides the template is the *medoid* conformation over training residues, which avoids the bond-length shrinkage that plain averaging would introduce. Trained distogram, pLDDT, and PAE heads provide confidence, with pLDDT targets computed on-the-fly as a stop-gradient soft lDDT. A post-hoc declash step (a short van der Waals-driven coordinate optimisation) roughly halves the clash rate at essentially unchanged lDDT.

#### A.6.2 Model training

QuickFold was trained from scratch on a single NVIDIA L40S GPU (46 GB) under a 480-minute wall-clock budget, reaching roughly 1.1 × 10^5^ optimisation steps (the exact count at the budget boundary is GPU-dependent, since the run is time-boxed rather than step-boxed). Optimisation used AdamW (learning rate 7 × 10^−4^, weight decay 10^−4^, gradient-norm clip 1.0) with an EMA of the weights (decay 0.999). The deployed weights are the EMA-averaged copy, which lowers run-to-run variance under the constant learning rate.

##### Scheduler and warmup

The learning rate follows a short 300-step linear warmup (0 → 7 × 10^−4^) and is then held constant with no decay. The warmup is the only step-based schedule, chosen to protect the first few hundred updates independent of hardware. Every other schedule is expressed as a *fraction of the wall-clock budget* rather than a step count, so a fixed-time run behaves consistently across GPUs. A crop curriculum grows the training crop from 96 to 144 to 192 residues at 0/45/70% of the budget (cheap short crops build local skill early, longer crops teach longer-range folding late), and the FAPE clamp is relaxed from 10 to 30 Å over the final 30% (local geometry first, medium-range gradients restored later).

##### Loss terms

The final objective has ten terms (Table 3). Six are active throughout; the four validity/global-fold terms (soft-TM, backbone continuity, backbone bond-angle, and van der Waals clash) are held at zero and linearly ramped in only over the last 40% of the budget, following the scheduling principle of SI.A.5 — geometric and superposition-based penalties destroy an unformed fold and become useful only once a rough fold exists.

**Table 3.** Final QuickFold loss terms. Ramped terms are held at zero until 60% of the wall-clock budget, then linearly increased to full weight by the end; all others are active throughout. Backbone terms apply only across genuinely sequence-adjacent residues.

| Term | Weight | Compares | Engages |
| --- | --- | --- | --- |
| FAPE | 1.0 | frame-aligned point error (6 rep. atoms/residue) | always |
| Distogram | 0.3 | C1'–C1' distance-bin cross-entropy | always |
| pLDDT | 0.1 | per-residue soft-IDDT confidence (stop-grad) | always |
| PAE | 0.1 | inter-frame aligned error (stop-grad) | always |
| $\chi$ | 1.0 | base-atom glycosidic-torsion coordinates | always |
| Soft-IDDT | 0.5 | sigmoid-relaxed IDDT on built C1' distances | always |
| Soft-TM | 1.0 | Kabsch-aligned soft TM-score on C1' | ramped ($\geq 60\%$) |
| Backbone continuity | 0.5 | O3'(i)–P(i+1) bond length | ramped ($\geq 60\%$) |
| Backbone bond-angle | 0.3 | phosphodiester junction bond angles | ramped ($\geq 60\%$) |
| Clash | 0.2 | vdW-aware inter-residue heavy-atom overlap | ramped ($\geq 60\%$) |

Figure 4 shows the training and validation loss curves for a representative seed.

**Figure 4.**
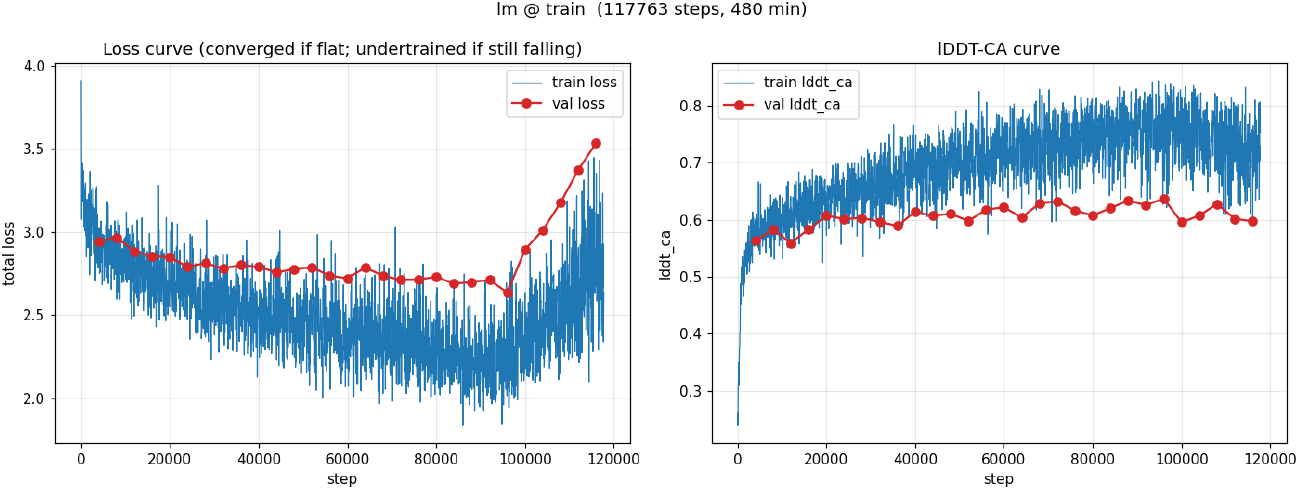
Training and validation loss curves for a representative QuickFold training run.

##### Checkpoint selection

Because the validity terms engage only in the last 40% of training, the global-argmax validation-lDDT snapshot is systematically the wrong choice: it lies in the pre-clash region and looks best on lDDT while being unusable (uncorrectable steric overlap). The deployable checkpoint is instead the best validation-lDDT snapshot among those in the last ~20% of training (step*/*step_max_ ≥ 0.80) whose clash rate has already settled, with ties broken by lower clash. The test split is never read during selection.

### A.7 Detailed model performance

#### Model size

QuickFold’s advantage in inference speed follows directly from a much smaller *trained* folding trunk (Table 4). QuickFold and RhoFold+ both carry a ~100 M-parameter frozen RNA language model, whereas NuFold uses none; the part that was developed and trained here is the 8.9 M-parameter trunk, 3× smaller than RhoFold+’s and 10× smaller than NuFold’s.

**Table 4.**
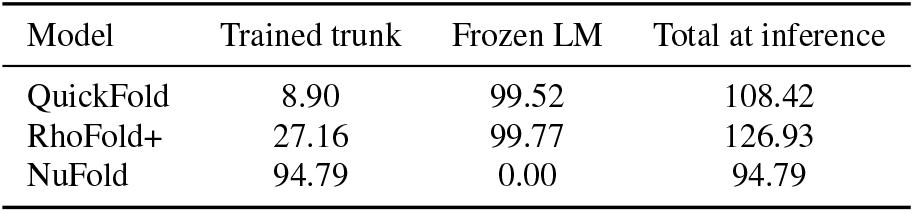
Parameter counts (millions). The trained trunk is the part developed in this work; the language model is frozen.

#### Full metrics

Table 5 extends the main benchmark with the global-alignment TM-score and RMSD from `barnaba` (RMSD in Å), and it reports both the raw model output and the minimized/declashed version of each tool (most-confident prediction, *n* = 80). Refinement changes the clash rate substantially but leaves the accuracy metrics almost unchanged, confirming that declashing QuickFold buys structural validity at negligible accuracy cost.

**Table 5.**
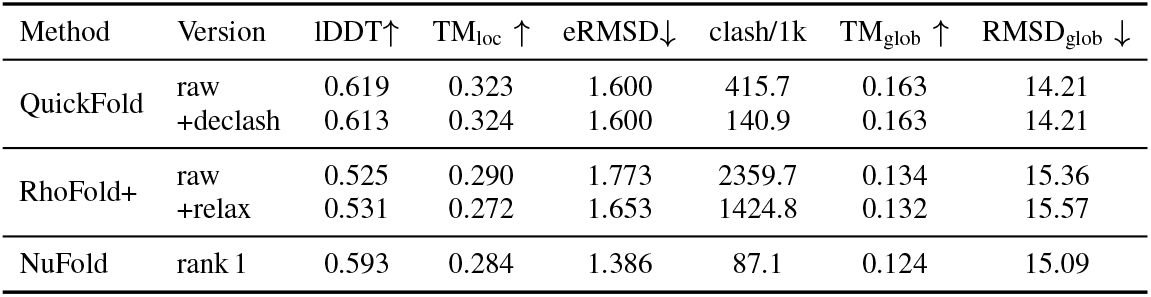
Extended test-set results (most-confident prediction, *n* = 80). “raw” is the model output; “+declash”/”+relax” are the refined versions. TM_loc_: USalign C3^*′*^ local alignment; TM_glob_/RMSD_glob_: barnaba global alignment;

#### Inference time

Table 6 breaks the per-target inference time into preprocessing (secondary-structure and language-model features), the folding-trunk forward pass, and refinement (MSA search excluded throughout). QuickFold’s folding trunk is 25× faster than RhoFold+’s and 215× faster than NuFold’s; its total is dominated by the optional declash step, and RhoFold+’s by Amber relaxation. This table does not consider MSA calculation, which tends to be the most expensive part of RhoFold+ and NuFold inference. QuickFold is MSA-free, so does not require this step.

**Table 6.** Per-target inference time (seconds), averaged over *n* = 80, MSA search excluded. RhoFold+ nests its language model inside the trunk, so preprocessing is not separately timed.

| Method | Preprocess (SS+LM) | Folding trunk | Refinement | Total |
| --- | --- | --- | --- | --- |
| QuickFold | 0.04 | 0.09 | 3.39 (declash) | 3.52 |
| RhoFold+ | — | 2.27 | 272.6 (Amber) | 274.9 |
| NuFold | 0.37 | 19.61 | — | 19.98 |

#### Deep-MSA subset

Most MSAs provided to RhoFold and NuFold were shallow MSAs. To better test the methods we assessed the performance on the subset of targets with alignment depth ≥128 (*n* = 15), QuickFold’s TM-score rises to ≈0.49 (Table 7). We caution that this slice is small, hence may not be representative.

**Table 7.** Deep-MSA subset (alignment depth ≥128, *n* = 15; most-confident prediction).

| Method | Version | IDDT↑ | $TM \uparrow$ | eRMSD↓ | clash/1k |
| --- | --- | --- | --- | --- | --- |
| QuickFold | raw | 0.639 | 0.489 | 1.645 | 771.1 |
|  | +declash | 0.628 | 0.489 | 1.641 | 232.4 |
| RhoFold+ | raw | 0.505 | 0.396 | 2.244 | 6633.0 |
|  | +relax | 0.494 | 0.379 | 2.269 | 6460.8 |
| NuFold | rank 1 | 0.551 | 0.304 | 1.468 | 97.4 |

#### A.7.1 Standalone inference protocol

All three models were run standalone over the same 80-target test set, with the model built or loaded once and looped over all targets, so per-target timing reflects compute only, not process startup or weight loading. RhoFold+ and NuFold were given the benchmark’s own precomputed A3M files (unmodified) as sequence/MSA input; no MSA database search was run for either, so their reported time covers feature preparation, the network forward pass, and each model’s own refinement, not database search. QuickFold never takes an MSA (SI.A.1).

- **QuickFold**. Three independently trained-from-scratch seeds (`seed0/1/2`), each a full training run; unlike NuFold’s ranks (below) these three predictions are genuinely independent samples. Input is single-sequence: a ViennaRNA base-pairing prior and a frozen RNA-FM embedding of the query. Refinement is QuickFold’s own coordinate-space declash (van der Waals-aware relaxation, 1500 steps, restraint weight 0.2). “Most confident” picks, per target, the seed with the highest mean pLDDT; “mean of 3” averages the metric over all three seeds.
- **RhoFold+**. Shipped pretrained weights, with RNA-FM nested inside the checkpoint. Refinement is its shipped Amber relaxation (OpenMM, 1000 steps, run on GPU). Each target’s relaxation ran in an isolated subprocess with a 900 s watchdog; 15/80 targets did not converge within that budget on our hardware and fall back to the unrelaxed prediction for the “+relax” rows. RhoFold+ is deterministic (*n*=1), so its “mean of 3” and “most confident” rows are identical.
- **NuFold**. Shipped pretrained weights (`global_step145245`). A per-target `ipknot` secondary structure is generated up front (timed separately). The forward pass uses the shipped defaults (`--repeat 1 --recycle 3`) and its own recycle-intermediate ranking; we kept `rank_1..rank_3`. NuFold ships no refinement. Its `rank_1..3` are recycle intermediates of a *single* forward pass, not independent samples, so their near-zero spread does not measure the same variance as QuickFold’s seed-to-seed spread.

### A.8 User feedback

#### How the human encouraged exploration

When the loop stalled, the human’s feedback was rarely a specific design. A representative prompt was:

> “don’t stop. try something new. Make bigger and bolder changes. think harder and try unconventional solutions. continue with the loop, use feedback from `Notes.md.`”

Roughly 60 of the 98 human turns were variants of this “be bolder” template. Concrete feedback fell into a few categories: scientific suggestions from visually inspecting predictions; bug reports from inspection; resource authorisation; and scope or objective decisions. Some substantive human turns are present in Table 8. The division of labour was consistent: the agent owned metrics, mechanisms, and implementation, while the human owned visual structural judgement, compute authorisation, and scope.

**Table 8.** Substantive human interruptions of the autonomous optimisation loop (turns that carried new information). Iteration ranges are the runs each turn set in motion

| Turn | Iter(s) | Contribution |
| --- | --- | --- |
| U11 | ~80 | New `experiments.tsv` columns broke the header schema; triggered the first ledger repair. |
| U16 | 126–130 | Directed a review of successful runs and an extended-budget study of how far converged trainings could go. |
| U17 | ~130 | Called the ledgers unusable; rebuilt them and requested the current best model and its architecture. |
| U18 | 131–154 | Set an execution plan, reverted to a 10–20 min budget for same-budget comparison, and mandated multi-metric reporting (TM-score, eRMSD, clash). |
| U34 | 200–205 | Reported missing validation PDBs and configs since iter197; triggered per-run `config.json` and reproducible eval outputs. |
| U48 | 229 | Select the best-performing validation epoch rather than the last, across all multi-checkpoint runs. |
| U49 | 229 | Evaluate *all* multi-checkpoint runs for a worth-saving checkpoint, not just one. |
| U50 | 202 | Audit caught models warm-started from a checkpoint; re-affirmed the train-from-scratch constraint (retired iter202). |
| U57 | ~260s | Endorsed the queue-many-runs mode, locking in batched overnight execution for the rest of the campaign. |
| U70 | 260 | Manually selected checkpoint step120000 (highest val IDDT, TM within noise) and placed a scope hold on new runs. |
| U89 | 289–294 | Proposed trying more RNA language models; broadened into a systematic 7-model sweep (RNA-FM confirmed optimal). |
| U93 | 279 (repro) | Diagnosed the epoch-selection rule as wrong (snapshot taken before declash engages) and commissioned a 3-seed reproducibility study. |
| U95 | 295–297 | Changed the objective to pursue an MSA-free variant, producing the final deployable model. |

## Supporting References

